# One Codex: A Sensitive and Accurate Data Platform for Genomic Microbial Identification

**DOI:** 10.1101/027607

**Authors:** Samuel S. Minot, Niklas Krumm, Nicholas B. Greenfield

## Abstract

High-throughput sequencing (HTS) is increasingly being used for broad applications of microbial characterization, such as microbial ecology, clinical diagnosis, and outbreak epidemiology. However, the analytical task of comparing short sequence reads against the known diversity of microbial life has proved to be computationally challenging. The One Codex data platform was created with the dual goals of analyzing microbial data against the largest possible collection of microbial reference genomes, as well as presenting those results in a format that is consumable by applied end-users. One Codex identifies microbial sequences using a *“k*-mer based” taxonomic classification algorithm through a web-based data platform, using a reference database that currently includes approximately 40,000 bacterial, viral, fungal, and protozoan genomes. In order to evaluate whether this classification method and associated database provided quantitatively different performance for microbial identification, we created a large and diverse evaluation dataset containing 50 million reads from 10,639 genomes, as well as sequences from six organisms novel species not be included in the reference databases of any of the tested classifiers. Quantitative evaluation of several published microbial detection methods shows that One Codex has the highest degree of sensitivity and specificity (AUC = 0.97, compared to 0.82-0.88 for other methods), both when detecting well-characterized species as well as newly sequenced, “taxonomically novel” organisms.

## 2 INTRODUCTION

As the efficiency, accuracy, and speed of high-throughput genomic sequencing (HTS) has continued to improve, a larger set of microbiologists working in clinical medicine, public health, and industry is adopting this powerful technology. Public health agencies are already beginning to use HTS to track the spread of outbreaks (Neimark 2015), and there is an increasing number of applications for which the exquisite precision and accuracy of genome-level identification justifies the (presently) higher marginal per-test cost. However, the broader adoption of HTS by scientists that may lack experience in bioinformatic analysis has created a need for bioinformatic solutions that are accessible for non-experts and provide high-quality analytical results (Grad 2014; Caboche 2014).

Taxonomic classification is the analytical basis of a wide range of applied microbiology supporting applications in public health, clinical diagnosis, and industrial production. Strain- and species-level taxonomic classification allows a user to identify specific pathogens, perform genomic epidemiology, and characterize microbial communities that may be associated with a particular phenotype. A variety of algorithms have been developed for such taxonomic classification, including the use of marker gene libraries (Segata 2012; Liu 2011), local alignment (Naccache 2014; Mitra 2011), and *k*-mer matching (Ames 2013; Wood 2014; Břinda 2015; Ounit 2015). *K*-mer-based analysis, the method used by Kraken, Seed-Kraken, CLARK, and One Codex, identifies short sequences (typically ranging from 17 - 31 bp) that are unique to specific taxa within a set of input reads. Based on the collection of *k*-mers that are found in a given read, it can be assigned to a particular taxon. By extension, a sample can be characterized according to the proportion of reads that are assigned to different taxa. Using this approach, microbial samples either from isolates or mixed samples can be characterized to the level needed to perform a large number of tasks needed for public health, clinical diagnosis, and industrial microbiology. As a public resource for academic data analysis, we believe it is valuable to provide the research community with a description of the performance and operation of the One Codex platform, while providing the raw data and analytical details needed to replicate or update such an evaluation as metagenomic methods continue to improve. In this paper we describe the functioning of One Codex and a rigorous functional evaluation of the state-of-the-art metagenomic classification methods, including the effect of database size on classification accuracy.

## 3 MATERIALS & METHODS

### 3.1 Taxonomic Classification Algorithm

One Codex classifies individual sequence reads according to the set of *k*-mers in that read that are unique to specific taxonomic groups. This analytical approach has been described extensively (Ames 2014; Wood 2014) and is implemented by One Codex using a default value of *k*=31. Briefly, each read is broken into the complete set of overlapping sequences of length 31bp that comprise it. These *k*-mers are compared against an exhaustive database that contains every *k-*mer and the taxonomic grouping to which it is unique (e.g., a specific clade of bacteria, archaea, or viruses). A compressed data structure is used to index and rapidly search *k-*mer databases generated from approximately 40,000 microbial genomes. Each read can then be summarized as a “*k-*mer hit chain” that describes the complete set of taxonomically-informative *k*-mers found in that read, as well as their positions. Individual reads are then assigned on the basis of the highest weighted taxonomic root-to-leaf path amongst these *k-*mer hits. For example, if a read has *k*-mers unique to *Enterobacteriaceae, Escherichia*, and *Escherichia coli,* it would be given the label *E. coli.* However, if it had *k*-mers unique to *Enterobacteriaceae, Escherichia*, and *Klebsiella*, it would be given the label *Enterobacteriaceae* – the most specific taxon that encompasses all detected *k*-mers, as *Klebsiella* and *Escherichia* are separate genera of *Enterobacteriaceae.* Finally, the distribution of reads from a single sample across different organisms and taxa is used to construct a comprehensive report that displays the organisms present in a sample.

### 3.2 Classification Accuracy

The goal of this evaluation effort was to assess the ability of a suite of bioinformatic methods to assign nucleotide sequences to the most accurate taxonomic group.

#### 3.2.1. One Codex Classification

Data were processed on the One Codex platform according to the instructions outlined in Section 2.3 – One Codex User Interface. One Codex uses two reference databases, the full One Codex database of approximately 40,000 bacteria, viruses, fungi, archaeal, and protists, and a smaller database containing the over 8,000 microbial genomes contained in the NCBI RefSeq database. Both the One Codex full database (referred to here as “One Codex”) and the One Codex RefSeq Database represent sequences available on July 8, 2015.

#### 3.2.2. Additional Classification Algorithms

The following metagenomic classification algorithms were downloaded, compiled, and installed according to the provided instructions in an Ubuntu environment on standard AWS EC2 instances (r3.8xlarge). In each case the indicated dependencies were installed as described and default run settings were used, except in the case of Clark in which the “RAM-light” flag was used in order not exceed system memory capacity.

- Metaphlan (2.1.0) - https://bitbucket.org/biobakery/metaphlan2
- GOTTCHA (1.0b) - https://github.com/poeli/GOTTCHA (database GOTTCHA_BACTERIA_c3514_k24_u2)
- Kraken (v0. 10.5-beta) - http://ccb.jhu.edu/software/kraken/
- Seed-Kraken (seedmod128b_from_0.10.6) - http://seed-kraken.readthedocs.org/
- Clark (v1.1.3) - http://clark.cs.ucr.edu/

#### 3.2.3. Additional Classification Databases

Metaphlan is distributed with a complete standalone database. Kraken is distributed with a reduced reference database (“Minikraken” - Dec. 8, 2014). In addition, we constructed the full Kraken database on July 5, 2015 according to the instructions provided at

http://ccb.ihu.edu/software/kraken/MANUAL.html. Given the contemporaneous snapshot of NCBI, the genomic content of the full Kraken database is roughly equivalent to that of the One Codex RefSeq Database, albeit with some differences in the exact repositories used. The Seed-Kraken database was constructed using the same complement of reference genomes as the Kraken database. The Clark bacterial reference database was constructed on Aug. 21, 2015 using the provided default instructions.

#### 3.2.4. Test Datasets

Two complementary approaches were employed to test the accuracy of these microbial detection methods. In the first, 500 sets of simulated reads (100,000 reads each) were generated from complete microbial genomes at a wide range of abundance levels in order to simulate a variety of biological assemblages (Mavromatis 2007). Each read set contained reads from 214 random genomes with roughly 100,000 reads simulated each, and additional reads from 10,425 genomes simulated at much lower abundance (1 – 30,000 reads each) (Figure 1). In total, 50 million reads were simulated from 10,639 genomes to provide a robust resource for the evaluation of metagenomic analysis methods. In every case, the genome was selected randomly from the set of complete genomes available in public sequence repositories, regardless of whether they were included in the One Codex database. Approximately 78% of the simulated genomes were also indexed in the One Codex Database, while 8.8% were indexed in the One Codex RefSeq Database. Importantly, the taxonomic ID for each read was encoded in the FASTQ header, allowing the direct comparison of the known source of each sequence against the taxonomic prediction made by each method.

**Figure 1.**
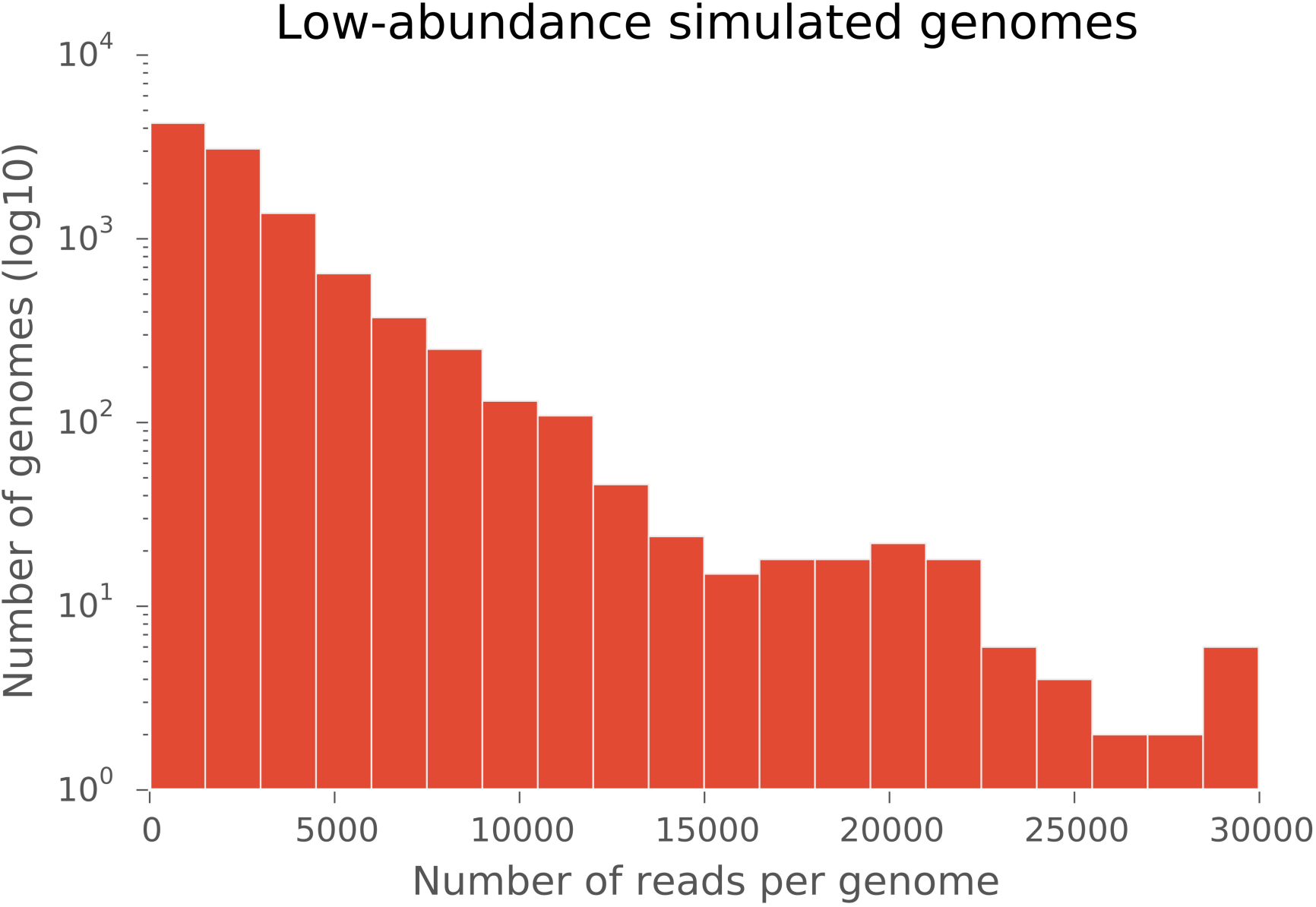
A frequency histogram of the number of reads simulated for the set of low-abundance genomes (10,425 genomes, 1 – 30,000 reads each). The horizontal axis shows the number of reads simulated per genome, and the vertical axis indicates the number of genomes simulated at that depth (log_10_). An additional set of 214 genomes were used to simulate at least 100,000 reads each.

In the second testing approach, a set of six organisms identified in the public repositories that were sequenced recently enough as to not be included in any of the reference databases. These “taxonomically novel” genomes did not have any other members of their species present in the reference database. Three of the six “taxonomically novel” test datasets were simulated from complete genome assemblies at 5X, two were raw Illumina reads, and one was an unassembled PacBio dataset (Table 1). While these datasets may contain contaminating or misidentified organisms, each analytical method will be challenged equally by those potentially confounding factors.

For both the single-isolate samples and the 500 sets of simulated reads (100,000 reads each), simulated 150bp single-ended reads were generated using the ART next-generation sequencing read simulator (v3.19.15) (Huang 2012) using the Illumina quality profile. For the single-isolate “taxonomically novel” test datasets, reads were generated at 5-fold coverage depth.

**Table 1.**
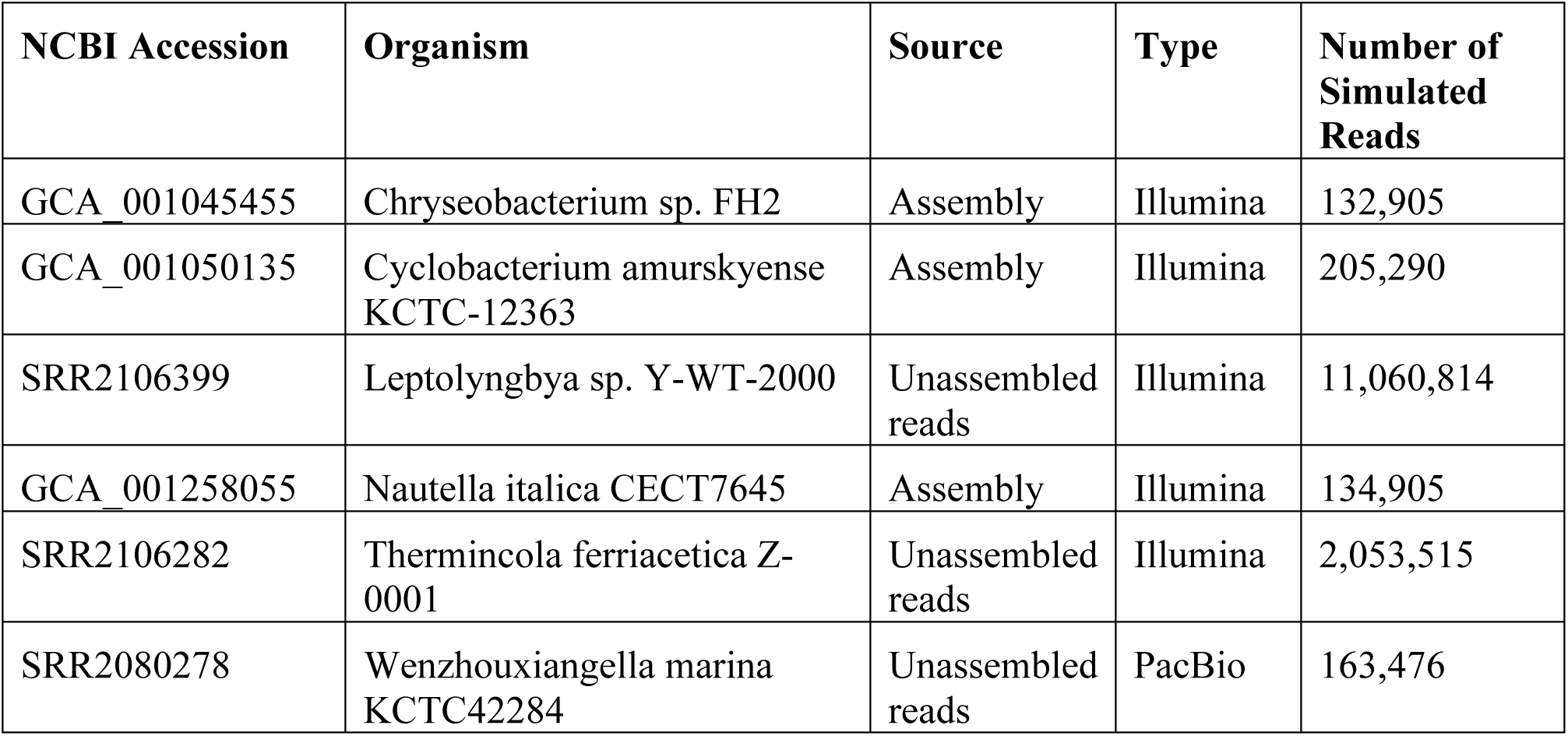
Datasets used to assess the performance of each method in identifying organisms not contained in the reference database.

| NCBI Accession | Organism | Source | Type | Number of Simulated Reads |
| --- | --- | --- | --- | --- |
| GCA_001045455 | Chryseobacterium sp. FH2 | Assembly | Illumina | 132,905 |
| GCA_001050135 | Cyclobacterium amurskyense KCTC-12363 | Assembly | Illumina | 205,290 |
| SRR2106399 | Leptolyngbya sp. Y-WT-2000 | Unassembled reads | Illumina | 11,060,814 |
| GCA_001258055 | Nautella italica CECT7645 | Assembly | Illumina | 134,905 |
| SRR2106282 | Thermincola ferriacetica Z-0001 | Unassembled reads | Illumina | 2,053,515 |
| SRR2080278 | Wenzhouxiangella marina KCTC42284 | Unassembled reads | PacBio | 163,476 |

#### 3.2.5. Statistical Summary

Kraken, Seed-Kraken, Clark, and One Codex provide taxonomic assignments for every read in a dataset, while Metaphlan and GOTTCHA provide an overall summary of dataset composition. Kraken, Seed-Kraken, Clark, and One Codex were evaluated with the complete set of 50 million simulated reads, while all methods were evaluated on the six single-organism “taxonomically novel” datasets. Each method was executed on an equivalent AWS EC2 instance (r3.8xlarge) with 12 processors available for parallelized steps.

For the set of 50 million simulated reads, the accuracy of classification by One Codex, Seed-Kraken and Kraken was evaluated on a read-by-read basis. One Codex was run with both the One Codex Database (~40,000 genomes) and the One Codex RefSeq Database (~8,000 genomes). Kraken was run with both the full database and the “Minikraken” database. Those methods assign an NCBI taxonomic identifier (‘taxid’) to each read. Given the known source of each read, the accuracy of the classification can be assessed across all levels of the taxonomy. For example, a read simulated from *E. coli* O157:H7 str. Sakai may be assigned to *Escherichia fergusonii*, in which case the species-level assignment is incorrect, while the genus-level assignment is correct (as is the family-, order-, class-, and phylum-). Similarly, a read simulated from *E. coli* O157:H7 str. Sakai may be assigned to *Enterobacteriaceae*, in which case the family-level assignment is correct and there is no assignment at the rank of genus, species, or strain.

Accuracy metrics were calculated as follows: Let A be the number of reads assigned correctly at a given taxonomic rank, B be the number of reads with any assignment at the given rank, C be the number of reads classified incorrectly at a less-specific rank, and D be the total number of reads. Sensitivity is defined as A / D, or the proportion of all reads assigned correctly at the given rank. Specificity is defined as A / (B + C), following Wood et al. (2015). For a set of reads simulated from *E. coli,* the classification of a read as *E. coli* would increase both species-level sensitivity and species-level specificity, classification as *Escherichia* would not increase species-level sensitivity, but it would increase species-level specificity, and classification as *E. fergusonii* would decrease both species-level sensitivity and species-level specificity.

Metaphlan and GOTTCHA do not assign taxa to individual reads, but rather predict the proportion of the dataset composed of different taxa. Therefore the specificity presented for those methods is the proportion assigned to the correct taxon at a given rank and no sensitivity metrics are reported.

### 3.3 One Codex User Interface

Samples are uploaded in FASTA or FASTQ format to the One Codex platform through a graphical upload tool with both drag-and-drop and folder navigation options. A command-line tool and API are also available for large-volume data upload (https://docs.onecodex.com). Once uploaded, reads are taxonomically classified and the interactive report is populated and linked to the user’s account (Supplemental Figures 1 and 2). The One Codex platform can be accessed at https://www.onecodex.com, and can be used freely to analyze public data.

### 3.4 Availability

Simulated datasets are available in a compressed FASTQ file containing 50M simulated reads at www.onecodex.com/data/papers/minot-krumm-greenfield-2015/simulated.reads.fastq.gz. The true taxonomic origin of each read is encoded in the FASTQ header as an NCBI taxid, allowing other researchers to replicate this analytical framework. One Codex is freely available for public use by academic researchers at https://www.onecodex.com. Supplemental Figures 1 and 2 show example screenshots of the One Codex platform.

## 4 RESULTS

### 4.1 Read-level Accuracy

We first summarized the accuracy of each tool on a per-read basis. One Codex showed the highest degree of sensitivity and specificity at each rank and the performance of the other methods varied with the database and assignment method used. It is notable that although the content of the Kraken and Seed-Kraken databases was identical, Seed-Kraken was more sensitive and less specific than Kraken at all taxonomic levels (Table 2 and Figure 2). While the reduced Minikraken database resulted in lower sensitivity and higher specificity than the full Kraken database, the reduced One Codex RefSeq Database was less accurate using both metrics. As noted above, ~78% of the simulated genomes were indexed in the One Codex Database, while 8.8% were indexed in the One Codex RefSeq Database. Accuracy metrics were also calculated for One Codex and One Codex RefSeq using only the subset of reads simulated from genomes not indexed in those databases. Species-level sensitivity and specificity for One Codex was 0.532 and 0.875, respectively, while One Codex RefSeq was 0.511 and 0.835. Similar comparisons could not be made for other methods without better characterization of the genome accessions used to create those reference indices.

**Table 2.**
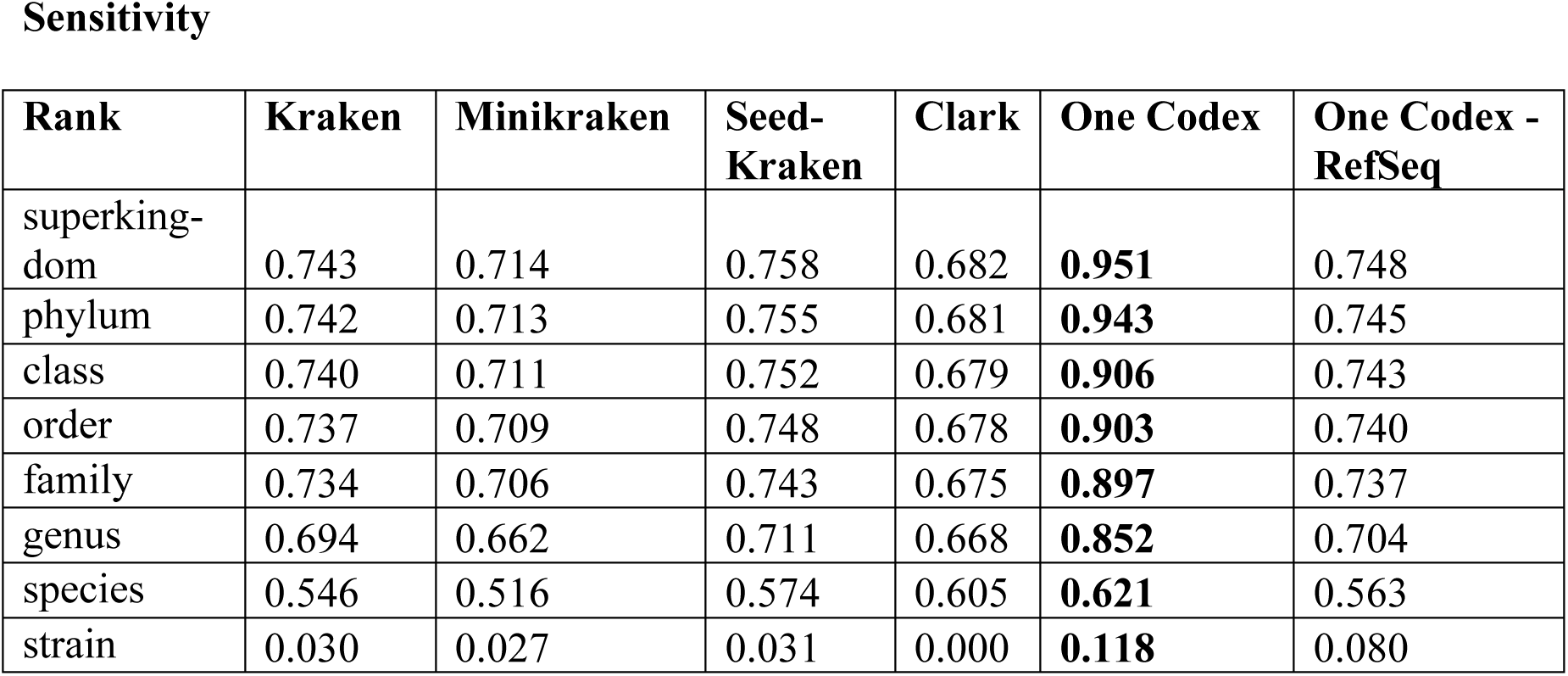
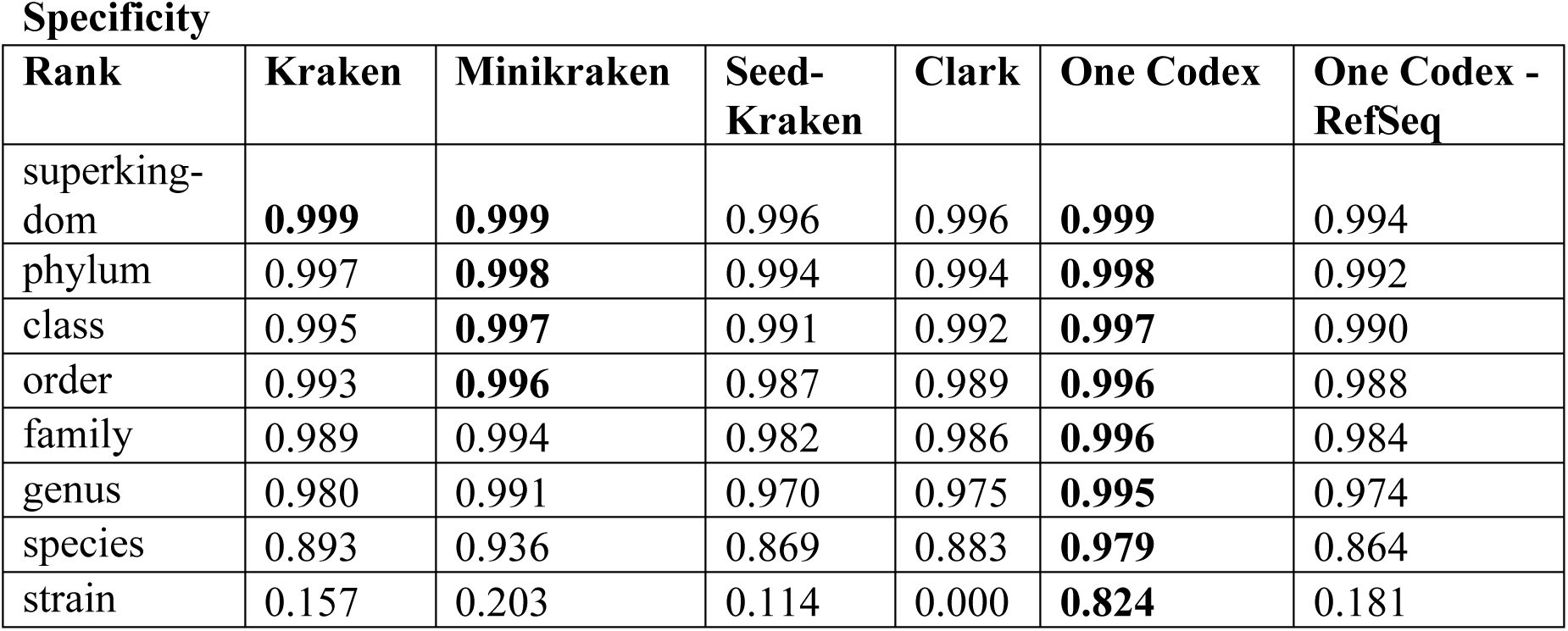
Summary of accuracy for six methods identifying the taxonomic origin of 50 million short sequence reads simulated from 10,639 microbial genomes. The maximum value at each rank is bolded.

Sensitivity
| Rank | Kraken | Minikraken | Seed-Kraken | Clark | One Codex | One Codex - RefSeq |
| --- | --- | --- | --- | --- | --- | --- |
| superkingdom | 0.743 | 0.714 | 0.758 | 0.682 | **0.951** | 0.748 |
| phylum | 0.742 | 0.713 | 0.755 | 0.681 | **0.943** | 0.745 |
| class | 0.740 | 0.711 | 0.752 | 0.679 | **0.906** | 0.743 |
| order | 0.737 | 0.709 | 0.748 | 0.678 | **0.903** | 0.740 |
| family | 0.734 | 0.706 | 0.743 | 0.675 | **0.897** | 0.737 |
| genus | 0.694 | 0.662 | 0.711 | 0.668 | **0.852** | 0.704 |
| species | 0.546 | 0.516 | 0.574 | 0.605 | **0.621** | 0.563 |
| strain | 0.030 | 0.027 | 0.031 | 0.000 | **0.118** | 0.080 |

| Rank | Kraken | Minikraken | Seed-Kraken | Clark | One Codex | One Codex - RefSeq |
| --- | --- | --- | --- | --- | --- | --- |
| superkingdom | **0.999** | **0.999** | 0.996 | 0.996 | **0.999** | 0.994 |
| phylum | 0.997 | **0.998** | 0.994 | 0.994 | **0.998** | 0.992 |
| class | 0.995 | **0.997** | 0.991 | 0.992 | **0.997** | 0.990 |
| order | 0.993 | **0.996** | 0.987 | 0.989 | **0.996** | 0.988 |
| family | 0.989 | 0.994 | 0.982 | 0.986 | **0.996** | 0.984 |
| genus | 0.980 | 0.991 | 0.970 | 0.975 | **0.995** | 0.974 |
| species | 0.893 | 0.936 | 0.869 | 0.883 | **0.979** | 0.864 |
| strain | 0.157 | 0.203 | 0.114 | 0.000 | **0.824** | 0.181 |

**Figure 2.**
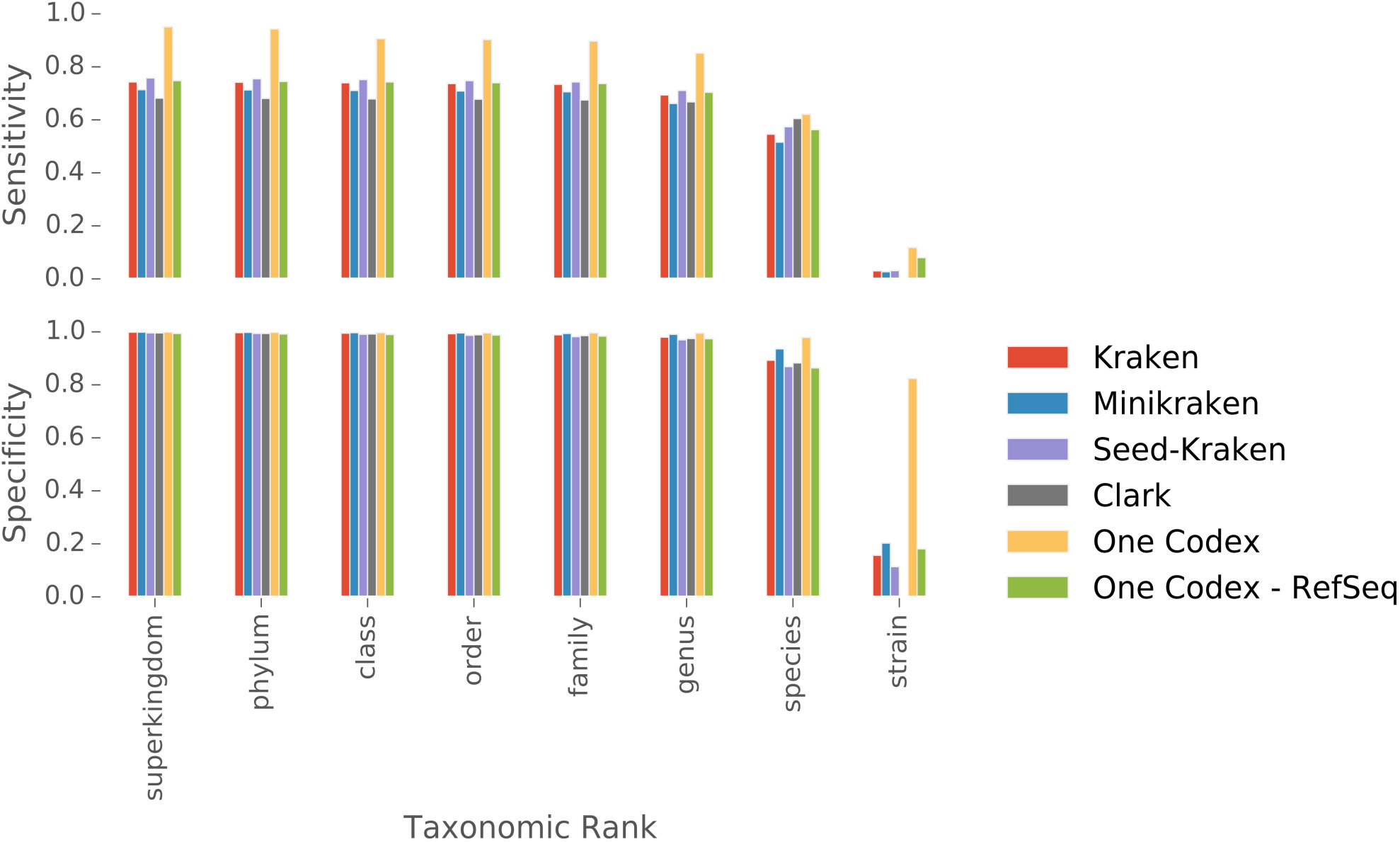
Summary of accuracy for six methods identifying the taxonomic origin of 50 million short sequence reads simulated from 10,639 microbial genomes.

### 4.2 Species-level presence/absence accuracy

We characterized the species-level accuracy of each classifier using a receiver operating characteristic (ROC) curve (Fig. 3). The ROC curve displays the true positive rate (TPR) and false positive rate (FPR) of species presence/absence across a range of read thresholds. Each dataset can be summarized as a set of species, each detected with a certain number of reads, and each marked as truly present, or truly absent. For any given number of reads, the species detected with at least that number of reads is marked as present, and any species with fewer than that number of reads is marked as absent. The TPR is calculated for a given read threshold as the number of true positive detections divided by the total number of total positives, and the FPR is calculated as the number of true negatives divided by the number of total negatives. The ROC curve for each method is shown in Figure 3.

**Figure 3.**
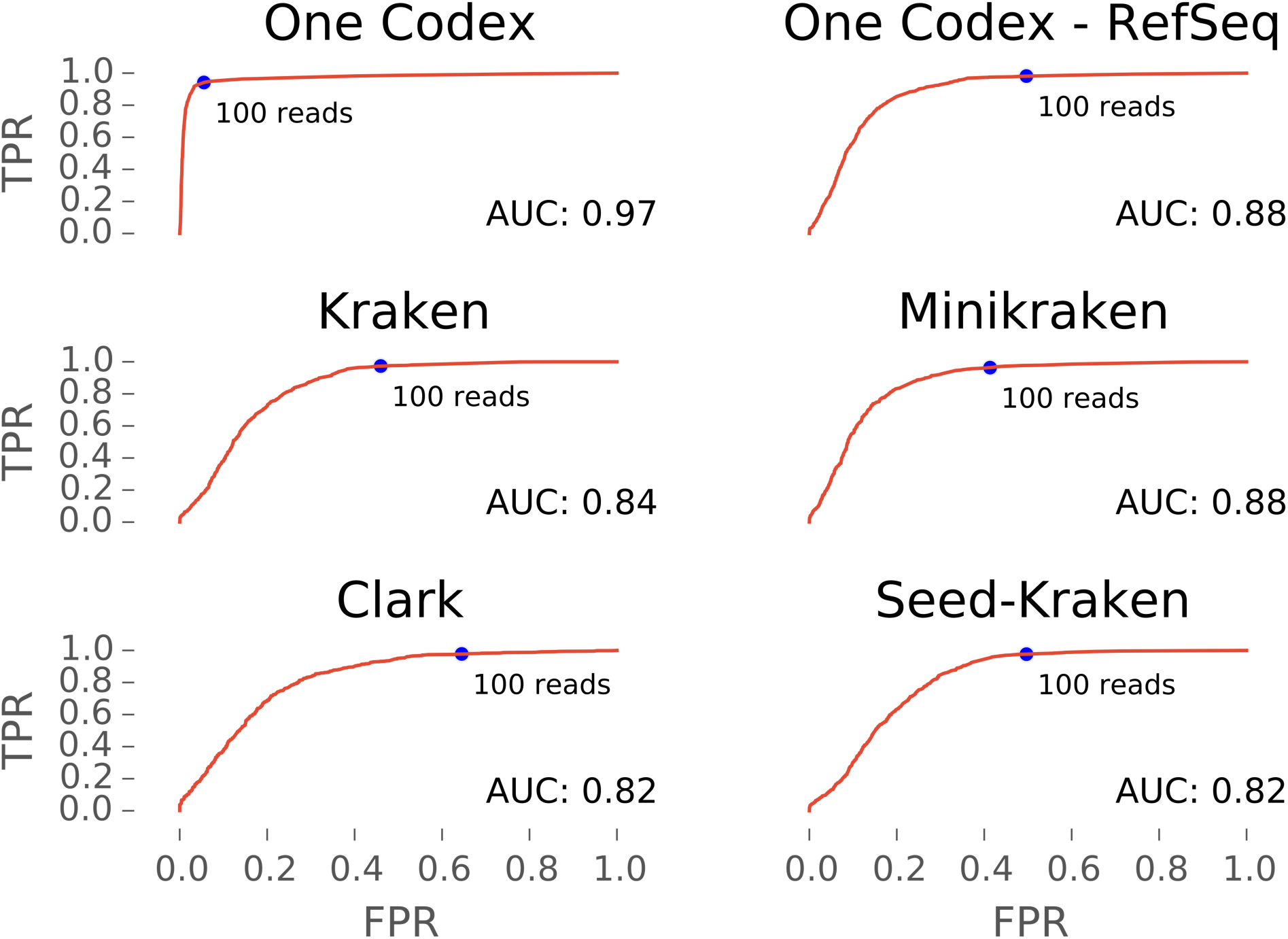
ROC curve displaying the performance of six taxonomic classification methods as specieslevel binary classifiers. Horizontal axis shows the False Positive Rate, and the vertical axis shows the True Positive Rate. Area Under Curve (AUC) is inset. The FPR and TPR are shown for a read cutoff value of 100 detected reads (150bp), which corresponds to roughly 0.003X coverage of a typical bacterial genome.

### 4.3 “Taxonomically Novel” Accuracy

Each of the “taxonomically novel” test datasets was selected because that species was not present in the reference database for any method. Due to the relative taxonomic novelty of these organisms, the closest match found by any method for these datasets was either at the genus-, family-, or order-level. For example, the most taxonomically similar reference organisms to *Wenzhouxiangella marina* KCTC 42284 (SRR2080278) share the order *Chromatiales*, as the family *Wenzhouxiangellaceae* was only proposed very recently (Wang 2015). Sensitivity and specificity metrics are shown in Table 3 alongside the rank at which the closest correct match was found by any method. GOTTCHA did not report any taxa above its threshold of detection for the three datasets showing ‘0’ in Table 3, and we provided the authors of that method with those datasets in order to confirm those results. Using the GOTTCHA ‘v20150825’ database they reported that dataset GCA_001045455 was assigned correctly at the family level and above, dataset SRR2106399 was assigned correctly at the order level, and dataset SRR2080278 did not have any assignments above the threshold of reporting (personal communication). Across all six datasets, One Codex displayed the highest sensitivity (0.391) while One Codex - RefSeq had the highest specificity (0.696).

**Table 3.** Accuracy of prediction for six organisms not found in any of the reference databases used by these methods. Note that Metaphlan and GOTTCHA results are presented as specificity metrics, as the abundance metrics reported by those methods are relative to the total classified composition of each sample (rather than the number of input reads).

| Dataset | GCA_001045455 | GCA_001050135 | SRR2106399 | GCA_001258055 | SRR2106282 | SRR2080278 |  |
| --- | --- | --- | --- | --- | --- | --- | --- |
| Assignment | Chryseobacterium | Cyclobacterium | Leptolyngbya | Rhodobacteraceae | Thermincola | Chromatiales |  |
| Rank | Genus | Genus | Genus | Family | Genus | Order |  |
| Sensitivity |  |  |  |  |  |  | Mean |
| Kraken | 0 | 0.149 | 0 | 0.203 | 0.831 | 0.005 | 0.198 |
| Minikraken | 0 | 0.096 | 0 | 0.093 | 0.787 | 0.001 | 0.163 |
| Seed-Kraken | 0 | 0.211 | 0.001 | 0.352 | 0.836 | 0.004 | 0.234 |
| Clark | 0 | 0.065 | 0.001 | 0.072 | 0.595 | 0.002 | 0.122 |
| One Codex | 0.313 | 0.225 | 0.01 | 0.965 | 0.827 | 0.005 | 0.391 |
| One Codex - RefSeq | 0.191 | 0.142 | 0.002 | 0.244 | 0.83 | 0.005 | 0.236 |
| Metaphlan | - | - | - | - | - | - | - |
| GOTTCHA | - | - | - | - | - | - | - |
| Specificity |  |  |  |  |  |  | Mean |
| Kraken | 0 | 0.961 | 0.013 | 0.920 | 0.995 | 0.102 | 0.499 |
| Minikraken | 0 | 0.985 | 0.04 | 0.943 | 0.998 | 0.266 | 0.539 |
| Seed-Kraken | 0 | 0.932 | 0.207 | 0.947 | 0.993 | 0.156 | 0.539 |
| Clark | 0 | 0.939 | 0.002 | 0.823 | 0.991 | 0.011 | 0.461 |
| One Codex | 0.920 | 0.927 | 0.109 | 0.999 | 0.988 | 0.082 | 0.671 |
| One Codex - RefSeq | 0.928 | 0.919 | 0.186 | 0.942 | 0.995 | 0.208 | 0.696 |
| Metaphlan | 0.935 | 1.000 | 0 | 1.000 | 0 | 0 | 0.489 |
| GOTTCHA | 0 | 1.000 | 0 | 1.000 | 0.971 | 0 | 0.495 |

### 4.4 Analysis Time

The time required for completing analysis of the 50M simulated reads for each method is presented in Table 4 (GOTTCHA and Metaphlan are not shown because they were not run on the read-level evaluation datasets). Although the complete set of 50M simulated reads were run in parallel batches of 10M reads, the time presented here is the cumulative processing time, rather than the shorter start-to-finish period of parallelized execution across multiple computational nodes. All methods were run with 12 processors on equivalent computational resources. Of these read-classification methods, Minikraken and Clark were the most rapid, and Seed-Kraken was the slowest.

**Table 4.** Computational time required for each method to classify the 50M simulated reads.

|  | Analysis Time (min:sec) |
| --- | --- |
| Clark | 04:50 |
| Minikraken | 05:15 |
| Kraken | 06:26 |
| One Codex | 22:13 |
| One Codex - RefSeq | 21:49 |
| Seed-Kraken | 45:58 |

## 5 DISCUSSION

The widespread adoption of high-throughput sequencing for the detailed characterization of mixed microbial samples presents an immense opportunity and challenge to the field of microbial genomics. Although genomic sequences can be used pinpoint the organisms in a sample down to the level of a single strain, accurate detection of each strain completely depends on the ability of a computational method to search for genomic sequences across the extent of known life. Not only does the volume of microbial reference genomes exceed that of the human genome by many-fold (181 billion bases of prokaryotic genome sequence can be found in NCBI as of Aug. 25, 2015), but the sequences found in wild-caught microbes may differ significantly from those of their domesticated relatives (Rinke, et al. 2013). In the face of these serious computational challenges, a large panel of computational methods have been proposed recently to perform the task of microbial detection (Oulas, 2015). However, it can be prohibitively difficult for a microbial researcher to rigorously evaluate all of the possible options in order to select the most appropriate method. To address that challenge, we have provided a comprehensive analysis of the performance of a wide range of the most widely adopted analytical methods. Moreover, we have made the test data and analytical framework available for others to evaluate future methods against a common reference. We believe that the large volume (over 50M simulated reads), phylogenetically complexity (10,639 randomly selected source organisms), and analytical portability (NCBI taxonomic identifiers recorded within read headers), makes this dataset a valuable resource for the research community.

One Codex provides the highest degree of accuracy, both sensitivity and specificity, across all taxonomic ranks, with 62.1% per-read sensitivity and 98.7% per-read specificity at the species-level (Table 2). The absolute performance of any detection method as a binary classifier was summarized by ROC analysis, showing that One Codex had the best performance (AUC: 0.97), with other methods performing roughly equally (AUC: 0.82 – 0.88). For purposes of illustration, the accuracy of each method was shown at an absolute abundance cutoff of 100 reads (Fig. 3). At that threshold for calling a species as present in a sample, One Codex showed a much lower FPR than other methods, suggesting that the larger set of reference organisms in the One Codex database serves to significantly reduce the number of species-level false positive detections with that method.

The array of methods evaluated here allows for an intriguing comparison of the effect of reference database and classification algorithm on overall performance. Kraken and Seed-Kraken use differing assignment algorithms and a common reference database, with Seed-Kraken providing higher sensitivity and Kraken providing higher specificity. This finding replicates similar precision/sensitivity trade-off that was observed previously for Seed-Kraken at the species-level (Břinda, 2015). While the performance of both Kraken and One Codex is presented with two alternate databases, the smaller Minikraken database was constructed by selecting a smaller number of kmers per organism, and the smaller One Codex RefSeq database was constructed by selecting a restricted number of reference organisms, so the resulting differences in sensitivity and specificity are not directly comparable. As the genomes encountered in real-world metagenomic samples rarely exactly match those found in reference databases, it is important to quantify the accuracy of detection for ‘out-of-reference’ genomes. Considering only the reads simulated from reads not found in those reference databases, the sensitivity and specificity of detection for One Codex was 0.532 and 0.875, respectively, while One Codex RefSeq was 0.511 and 0.835, reflecting only a modest decline in performance for that large group of ‘out-of-reference’ genomes. Further research into the effect of reference database composition on predictive accuracy could conceivably enable the creation of classification methods with smaller computational footprints and improved performance.

The largest difference in performance between classification methods can be seen in the “taxonomically novel” test datasets. Each method detects a different subset of organisms, indicating that the composition of each reference database highly determines the ability of a method to detect a given organism. Overall, One Codex had the highest average sensitivity (39.1%), which was far higher than the next-most sensitive methods One Codex RefSeq (23.6%) and Seed-Kraken (23.4%). This set of six datasets is a useful demonstration of the fundamental challenge of accurately classifying sequences from organisms not found in any reference database. Even when the closest possible rank is at the genus or above, each method varies widely in its ability to assign sequences correctly to that rank. The highly sensitive performance of One Codex against these samples suggests that a large and comprehensive reference database not only enables more accurate detection of well-characterized taxa, but also enables more accurate detection of taxonomically-novel and phylogenetically divergent organisms.

By evaluating a wide range of taxonomic classification algorithms against a large and complex set of 10,639 simulated genomes, as well as a set of six recently sequenced and phylogenetically-distinct organisms, we have generated important insight into the ability of microbial researchers to accurately characterize unknown metagenomic samples. Most notably, because Kraken, Minikraken, One Codex, and One Codex RefSeq all classify reads using taxonomically-unique 31mers, the differing performance of these methods is undoubtedly due to the much different composition of those databases, with larger reference databases leading to greater analytical accuracy. These results show the value of continually expanding reference database collections in order to more accurately classify the vast pool of unknown, unsequenced microbial ‘dark’ matter (Rinke 2013), as well as the specific strains of well-known pathogens that cause human disease.

## 6 ACKNOWLEDGEMENTS

*Conflict of Interest:* Authors are employed by Reference Genomics, Inc., which develops the One Codex platform.

## 8 SUPPLEMENTAL FIGURES

**Supplemental Figure 1.**
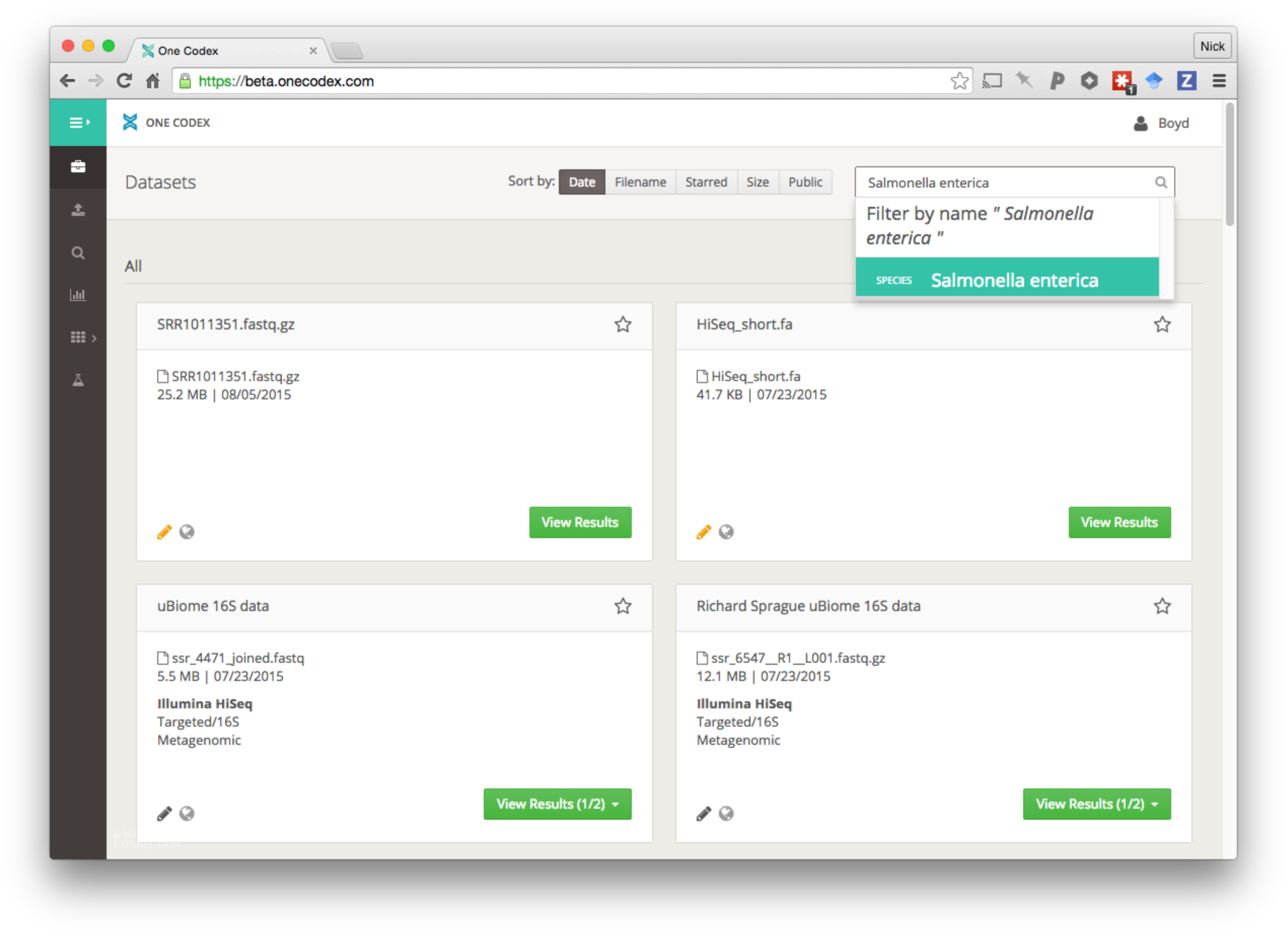
Example of dataset browser page on the One Codex platform. Datasets are organized by metadata (e.g. name, date, size, environment, platform, etc.) and user-defined tags. Figure displays an example of the user searching for datasets containing *Salmonella enterica* above a specified abundance threshold.

**Supplemental Figure 2.**
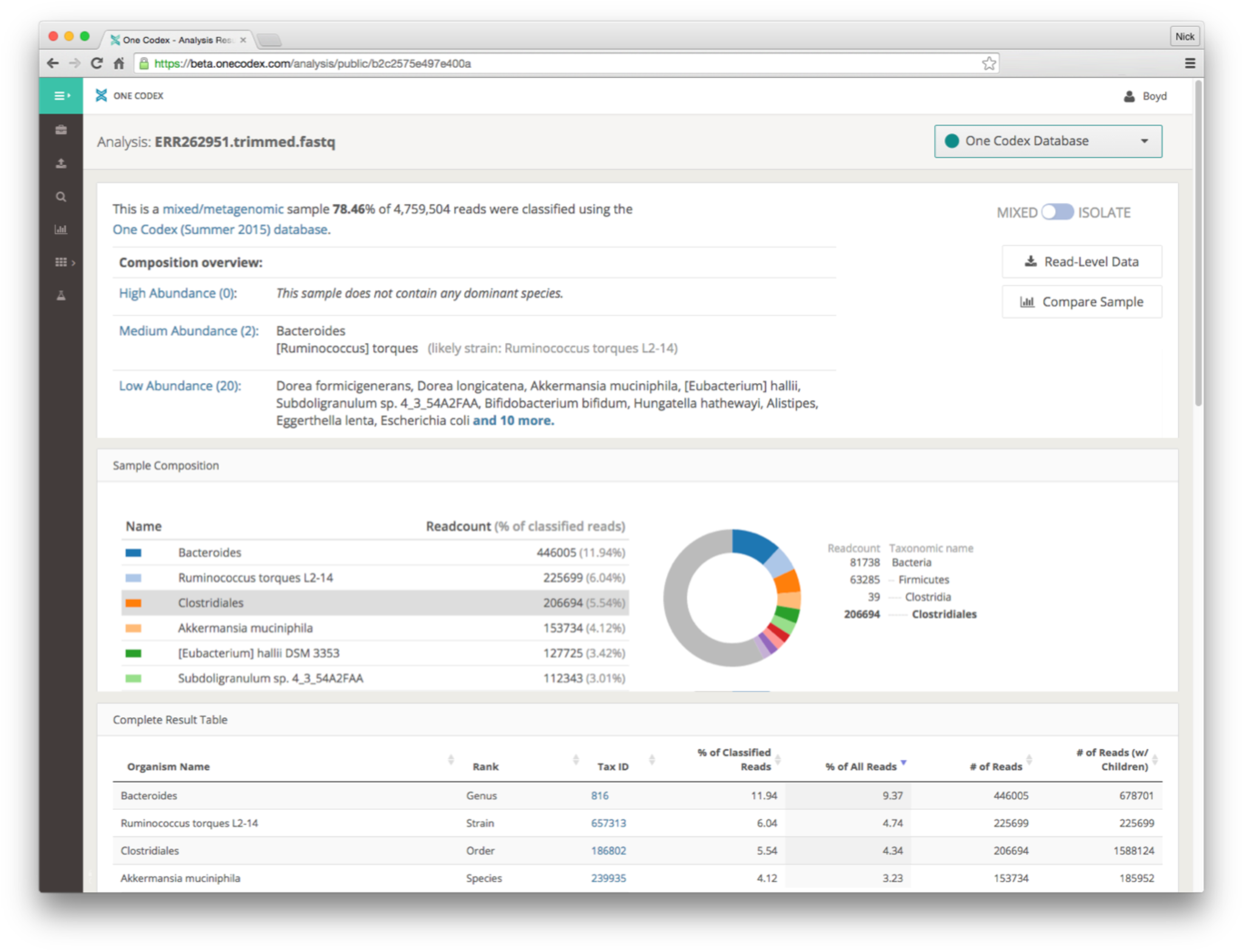
Example of metagenomic analysis display for a single sample. Users have the option of downloading read-level assignments or dataset summaries, comparing the abundance profile against that of other samples, and navigating to additional analyses for each dataset.

